# Lamin A/C safeguards replication initiation by orchestrating chromatin accessibility and PCNA recruitment

**DOI:** 10.1101/2025.03.06.641818

**Authors:** Mengling Zhang, Wenxue Zhao, Zhiguang Xiao, Lei Chang, Xiaotian Wang, Yichen Bai, Xiaodong Guan, Tianyan Liu, Yongzheng Li, Qing Li, Cheng Li, Dongyi Xu, Qian Peter Su, Yujie Sun

**Author notes:** These authors contributed equally to this work (Mengling Zhang, Wenxue Zhao, Zhiguang Xiao).

## Abstract

Lamin A/C, a nuclear lamina protein, is essential for maintaining nuclear architecture, organizing chromatin and preserving genomic stability. However, its role in regulating DNA replication remains unclear. This study investigates how Lamin A/C regulates proper activation of replication origins by orchestrating the chromatin structure and recruitment of proliferating cell nuclear antigen (PCNA). We demonstrate that Lamin A/C stabilizes replication domains (RDs) by restricting chromatin mobility, preserving spatial organization, and maintaining accessibility. Furthermore, Lamin A/C interacts with PCNA and sequesters a pool of PCNA, to regulate its recruitment to replication machinery. The loss of Lamin A/C leads to an increase in chromatin dynamics, RD accessibility, and PCNA availability at RDs, which together trigger excessive activation of replication origins, leading to replication stress and DNA damage. These disruptions prolong the S phase and compromise genome stability, highlighting Lamin A/C as a critical gatekeeper of balanced replication initiation. Our findings reveal Lamin A/C’s dual role in chromatin organization and replication machinery regulation, offering valuable insights into its involvement in replication-associated diseases and potential therapeutic opportunities through targeting replication dynamics.

## Introduction

The precise regulation of DNA replication initiation is essential for maintaining genomic stability. Errors during replication can lead to replication stress, DNA damage, and genomic instability—hallmarks of cancer and other replication-associated diseases^1,2^. In mammalian cells, the replication of chromatin DNA begins at tens of thousands of potential replication origins. DNA replication initiation is tightly regulated by a two-step process: origin licensing and origin firing, which together ensure the accurate duplication of the genome in both spatial and temporal dimensions^3–7^. During origin licensing, initiator proteins, including the Origin Recognition Complex (ORC) and the Minichromosome Maintenance (MCM) complex, are sequentially assembled onto these potential origins, designating them for future activation. This is followed by origin firing, where additional factors such as the Cdc45-MCM-GINS (CMG) helicase complex are recruited to a small subset (∼10%) of these licensed origins, triggering the formation of bidirectional replication forks^8,9^. Origin firing then marks the onset of replication, driven by the assembly of the replisome machinery, including Proliferating Cell Nuclear Antigen (PCNA), a key factor for DNA synthesis and elongation^10,11^. The delicate regulation of this process is crucial, as both under- and over-activation of replication origins can disrupt the balance necessary for proper DNA replication. Insufficient activation of replication origins may compromise the completion of genome replication, while excessive origin activation can disrupt replication fidelity by inducing replication stress^12–16^. The complexity of replication regulation in higher eukaryotes is likely a consequence of their larger genomes and more intricate chromatin organization, highlighting the need for robust spatiotemporal coordination to ensure genome stability.

Extensive studies have shown that DNA replication initiation is closely coordinated with the spatiotemporal organization of chromatin^17–20^. In metazoan cells, replication origins are not randomly distributed, but are clustered within megabase (Mb)-sized chromosomal regions known as replication domains (RDs), which serve as functional units for coordinating replication initiation^1,9^. Each RD encompasses multiple replication origins, whose spatial positioning and activation patterns are governed by factors such as chromatin accessibility, nuclear organization, and the local epigenetic landscape^21^. These domains replicate in a defined temporal order during the S phase, with their replication timing correlating with the chromatin architecture. Chromatin architecture, defined by the hierarchical folding of the genome, consists of various structural elements, including nucleosomes, topologically associating domains (TADs), and higher-order compartments^22–25^. At the compartment level, transcriptionally active A compartments generally replicate earlier, while transcriptionally inactive B compartments replicate later^23,26^. TADs, which are conserved structural units of the genome, spanning ∼0.5–1 Mb, help organize chromatin into functional domains^24,27^. The alignment of RD boundaries with TADs suggests that these chromatin structures function as units that coordinate replication^28^. Advances in high-resolution imaging techniques, such as Structured Illumination Microscopy (SIM)^29^, STimulated Emission Depletion microscopy (STED)^30^, STochastic Optical Reconstruction Microscopy (STORM)^31–34^ and Expansion Microscopy (ExM)^35,36^, have provided unprecedented insights into the spatial organization of chromatin and its impact on DNA replication. These cutting-edge technologies allow for the precise visualization of RDs and their spatial organization at the nanometer scale, thereby enabling detailed investigations into how chromatin structure governs the spatiotemporal regulation of replication initiation.

In addition to chromatin architecture, nuclear scaffold proteins have also been suggested to play an important role in regulating DNA replication^37–40^. Lamin A/C, a key component of the nuclear lamina, maintains nuclear integrity and regulates chromatin dynamics^41–44^. Unlike B-type Lamins, which preferentially remain anchored to the nuclear envelope due to permanent farnesylation, Lamin A/C undergoes a transient farnesylation followed by proteolytic cleavage, resulting in its mature form^45–48^. Matured Lamin A/C primarily localizes to the nuclear periphery, while a relatively dynamic and soluble pool remains in the nuclear interior^49^. Lamin A/C’s unique dual distribution enables it to support nuclear architecture while simultaneously regulating chromatin structure and nuclear processes, with its deficiency being associated with increased chromatin dynamics^50,51^, altered chromatin structure^52,53^, replication stress^47,54,55^, and replication-related diseases such as cancer and HPV infection^46,56^. Beyond its structural role in chromatin regulation, Lamin A/C also interacts with PCNA^57,58^, implying a potential regulatory role in replication. Yet, the precise mechanism by which Lamin A/C regulates DNA replication remains unclear, particularly whether its influence is exerted directly or mediated through chromatin organization. Our previous study presented a distinct chromatin-regulated spatial pattern of replication initiation and propagation, revealing that PCNA remains localized at the periphery of RDs from the G1 phase to the G1/S transition^32,33^ This positioning appears critical for selective initiation of replication origins, potentially reflecting an interplay between PCNA and nuclear scaffold proteins such as Lamin A/C. However, whether Lamin A/C facilitates PCNA recruitment to RDs during replication initiation remains unclear, leaving the precise role of the Lamin A/C-PCNA interaction in DNA replication to be further elucidated.

In this study, we investigate the multifaceted role of Lamin A/C in regulating DNA replication initiation and maintaining genome stability. Utilizing advanced imaging methods combined with sequencing techniques, we demonstrate that Lamin A/C acts as a gatekeeper, restricting the number of active replication origins. Our findings show that Lamin A/C deficiency alters the organization of replication domains, increases PCNA recruitment to RDs, and results in excessive activation of replication origins, which leads to replication stress and increased DNA damage. To explain these findings, we propose a model in which Lamin A/C simultaneously stabilizes the spatial structure of replication domains and modulates the availability of PCNA, thereby maintaining a balanced replication initiation program and preventing excessive origin firing. By elucidating Lamin A/C’s dual role in chromatin organization and replication machinery regulation, this study provides new insights into its critical function in preserving genome integrity.

## Results

### Lamin A/C Deficiency Enhances DNA Replication Origin Firing in Early S Phase

To investigate the functional impact of Lamin A/C deficiency, we utilized CRISPR knock-out to generate Lamin A/C-deficient cell lines (Hela S3 and MD-MBA-231). RNA-seq, combined with Kyoto Encyclopedia of Genes and Genomes (KEGG) pathway enrichment analysis, revealed that the loss of Lamin A/C led to differential expression of genes significantly enriched in pathways related to cancer HPV infection, both of which are associated with DNA replication (**Supplementary Fig. 1a, b**). Additionally, Gene Ontology (GO) enrichment analysis showed that these differentially expressed genes were enriched in biological processes such as the cell cycle, cell proliferation, and DNA damage (**Fig. 1a; Supplementary Fig. 1c**). Specifically, E2F2 transcription factor (**Supplementary Table 1),** which are key regulators of DNA replication initiation and directly influence PCNA expression (discussed below), were enriched in these biological processes. Other key replication-related genes, such as replication licensing factor CDC6 and MCM8 proteins, were also enriched (**Supplementary Table 1**). These findings underscore Lamin A/C’s pivotal role in orchestrating DNA replication and maintaining genomic integrity.

**Figure 1.**
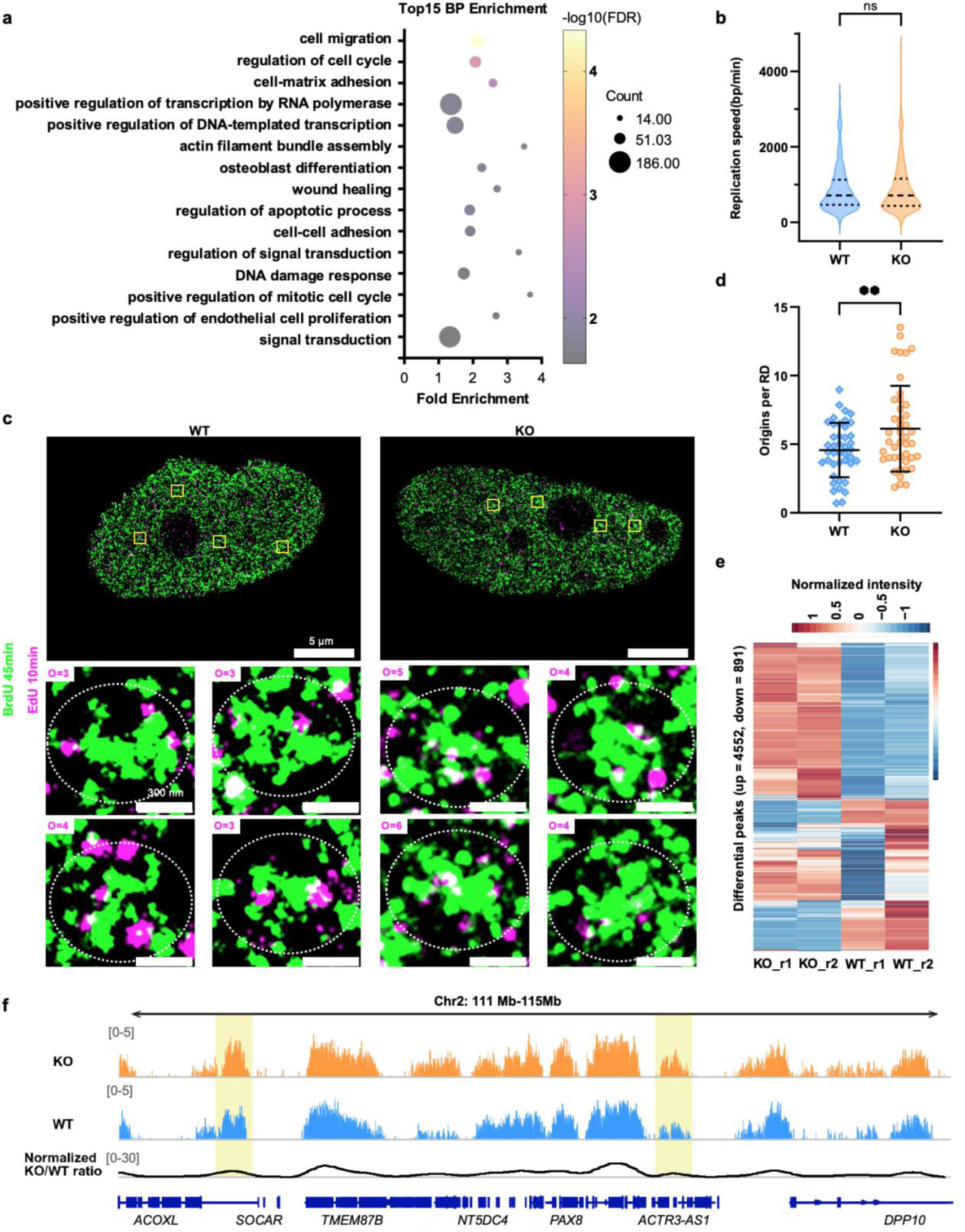
Lamin A/C depletion leads to increased replication initiation. **(a)** Top 15 enriched GO biological process (BP) terms in HeLa Cells. The bubble plot shows significantly enriched biological processes, with fold enrichment on the x-axis, bubble size representing gene count, and color indicating -log10(FDR). **(b)** Violin plot comparing replication fork speed between WT and KO cells. Data are shown as median ± quartiles. Significance levels are indicated as follows: ns, P>0.05. **(c)** Representative STORM images showing the distribution of replication origins and replication domains (RDs) in WT (left) and Lamin A/C knockout (KO, right) cells. Cells were synchronized, and RDs were labeled during the early S-phase with a 45-min BrdU pulse. A second synchronization was performed, and origins were labeled during the early S-phase with a 10-min EdU pulse. Scale bars, 5 μm. Enlarged rendered views of the yellow-boxed regions are shown below. Scale bars, 300 nm. The calculated number of origins within each targeted RDs (white circles) is indicated in the top-left corner of each magnified view. **(d)** Quantification of replication origin number per RD in WT and KO cells from the experiments shown in (c). Data are presented as mean ± SE, with n ≥ 30 cells. Significance levels are indicated as follows: ** P<0.01. **(e)** Heatmap of differentially regulated BrdU-seq peaks (>10 Kb). Heatmap displaying differentially regulated peaks of early S-phase 10-min BrdU labeling, representing replication initiation signals, comparing WT and KO cells. Red indicates upregulated signals, while blue indicates downregulated signals. Data are derived from two independent replicates. **(f)** The representative genomic region of BrdU-seq signals. Representative genomic region (hg38, Chr2: 111 Mb – 115 Mb) showing changes in BrdU-seq signals between WT (middle, blue) and KO (top, orange) samples. The bottom panel displays the normalized ratio (KO/WT) to highlight differential replication timing. The yellow shaded regions indicate representative peaks exhibiting changes. Gene annotations for this region are shown below for reference.

To study how Lamin A/C influences replication initiation, we synchronized cells to the G1/S boundary, (Methods; **Supplementary Fig. 2a**), and then imaged the replication initiation signals by 10-min pulse labeling of the replication as cells released into the S phase^33,59^. In wild-type cells, early replication signals were observed as the characteristic firing "foci". Interestingly, under Lamin A/C deficiency, these replication initiation signals were significantly enhanced. This phenomenon was consistent across both HeLa and 231 cell lines (**Supplementary Fig. 2b-e**). Besides, no replication signals were detected before S phase release in either wild-type or Lamin A/C deficient cells (**Supplementary Fig. 2c**), ruling out the possible effect of Lamin A/C loss on cell synchronization.

Quantitative analysis indicated that replication foci, numbering around 100 per cell (**Supplementary Fig. 2f, g**), possess an average radius of ∼300 nanometers (**Supplementary Fig. 2h, i**). Notably, the fluorescence intensity of individual replication foci was significantly increased in Lamin A/C deficient cells (**Supplementary Fig. 2j, k**). As each individual replication focus contains multiple replication origins, the increased initiation signal could be resulted from either more activated replication origins or an increase in replication fork speed. Using DNA fiber assays, we measured the replication fork speed and found it unchanged between Lamin A/C-deficient cells and wild-type cells (**Fig. 1b**). These findings exclude the possibility that changes in replication fork speed accounted for the upregulation of replication initiation.

To further determine whether the increase in replication initiation signals arose from an increase in activated replication origins, we performed dual-labeling of replication domains and initiation signals, followed by STORM super-resolution microscopy with ∼20 nm resolution^31^. The 10-min pulse-labeled foci precisely mark replication initiation sites, as EdU was added immediately after replication arrest release, and their ∼30 nm size, revealed by super-resolution imaging, aligns with STORM’s spatial resolution, ensuring accurate localization^33^. Building on our prior findings^33^, these results confirmed the peripheral localization of replication origins in wild-type cells and also revealed that this spatial peripheral distribution is preserved even in the absence of Lamin A/C (**Fig. 1c**). Notably, the number of replication origins per RD increased by approximately 20% in Lamin A/C-deficient cells (**Fig. 1c, d**). Together, these data show that the loss of Lamin A/C results in an increase of activated replication origins.

To validate these observations, we used BrdU-seq to measure replication initiation activity in wild-type and Lamin A/C deficient cells during early 10-min S phase entry. Consistent with the imaging results, 83% of the 5,443 replication peaks demonstrated increased peak heights in Lamin A/C knockout cells compared to wild-type cells (**Fig. 1e; Supplementary Fig. 3a**). Notably, Lamin A/C deficiency did not alter the locations of replication initiation peaks, which remained consistent with those in wild-type cells and were predominantly confined to early replication domains within A compartments, with no new peaks observed in middle or late replication domains (**Fig. 1f; Supplementary Fig. 3b**). Given that each peak by 10-min BrdU-seq typically spans around 30-100 kb (**Supplementary Fig. 3c**), these peaks represent clusters of multiple replication origins. This upregulation of replication peaks aligns with the increased number of replication origins observed through microscopy (**Fig. 1c, d**). Additionally, upregulated replication peaks were more likely to be located near TAD boundaries compared to downregulated peaks (**Supplementary Fig. 3d, e**). These results confirm that the effect on early replication initiation by Lamin A/C deficiency is primarily due to an increase in the number of replication origins.

Moreover, to rule out potential replication stress introduced by the aphidicolin-involved synchronization method, which might activate dormant replication origins^60,61^, we employed a replication stress-free method (Methods; **Supplementary Fig. 4a**) to observe the replication initiation process from the G1 to S phase. Notably, after early S phase entry (**Supplementary Fig. 4b**), replication signals in Lamin A/C-deficient cells rapidly increased, significantly higher than in wild-type cells. This finding is consistent with results from synchronization experiments, and adds further support to the reliability of replication initiation signal detection under synchronized conditions.

### Loss of Lamin A/C Increases Replication-induced DNA Damage and Genome Instability

Replication initiation is a tightly regulated process crucial for maintaining genomic stability. Both insufficient and excessive replication origins can disrupt this balance, leading to replication stress^13,16^. To investigate whether unscheduled replication initiation leads to replication stress under Lamin A/C deficiency, we analyzed γH2AX expression levels before and after replication initiation in G1 and S phases via Western blotting. The results showed that DNA damage, as indicated by increased γH2AX levels, was more pronounced in the S phase than in the G1 phase, both in wild-type and Lamin A/C-deficient cells (**Fig. 2a, b; Supplementary Fig. 5a, b**). Additionally, we conducted dual-color super-resolution imaging to label replication origins and replication stress markers. Further analysis revealed that γH2AX sites were consistently located near replication origins under both normal and Lamin A/C-deficient conditions, with a statistically significant proximity compared to the random distribution control (**Fig. 2c, d**). This average distance of approximately 180 nm—slightly smaller than the typical size of an RD (calculated below)—suggests that replication stress and initiation events are spatially correlated at the level of RDs. These findings suggest that the increased number of replication origins observed under Lamin A/C deficiency enhances the likelihood of replication stress, highlighting a critical link between active origin density and genomic stability.

**Figure 2.**
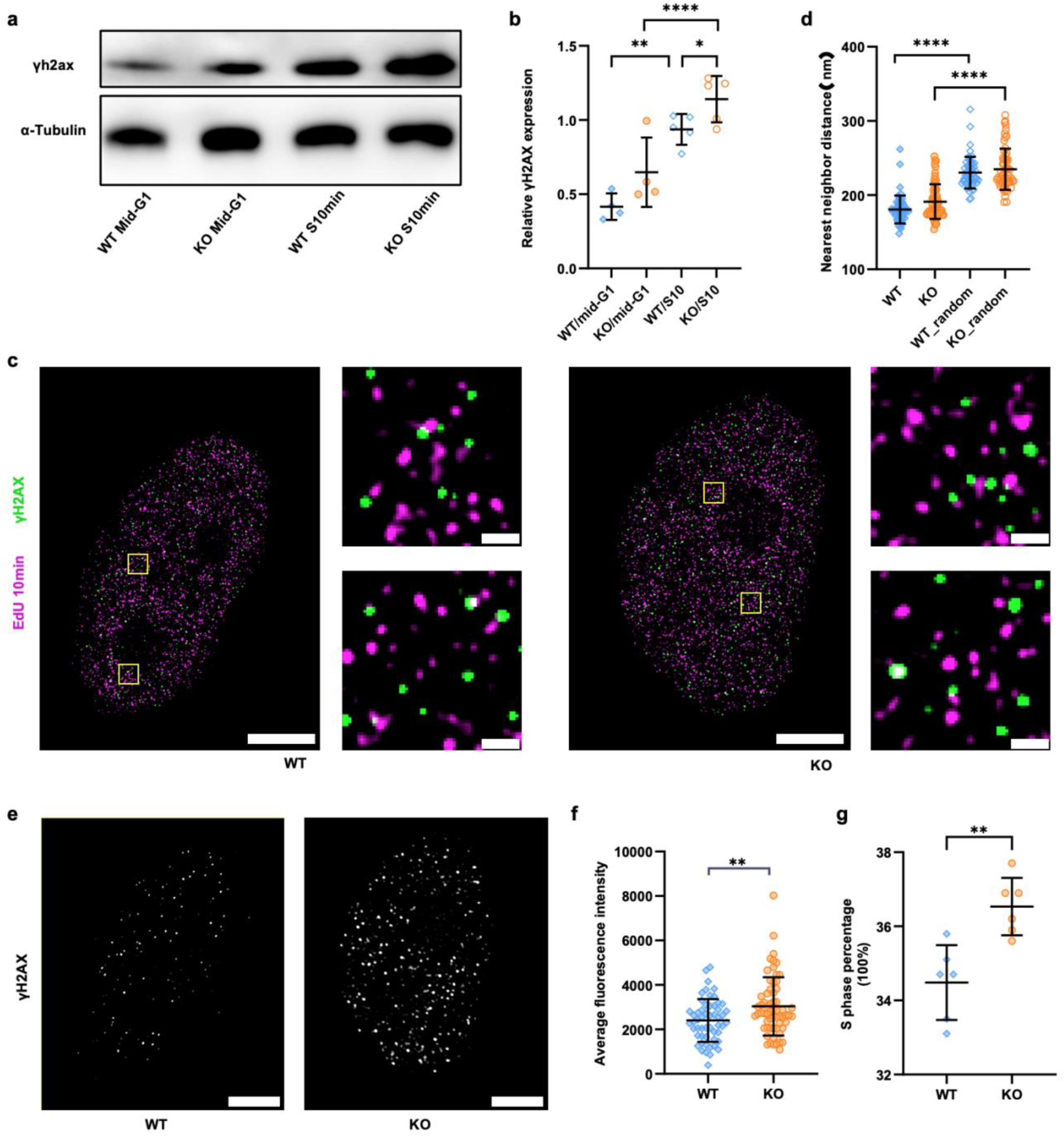
Replication damage in WT and Lamin A/C knockout cells. **(a)** Immunoblot analysis of γH2AX in WT and KO cells during mid-G1 phase (5 hours after release) and after 10 minutes of early S-phase. Tubulin is used as a normalization control. **(b)** Quantification of γH2AX signals from the experiments shown in (a). n ≥ 30 cells. **(c)** Representative STED images showing the spatial distribution of γH2AX (green) and replication initiation signals in WT (left) and Lamin A/C knockout (KO, right) cells. Replication initiation signals are labeled with EdU (magenta) after 10 minutes of early S-phase. Scale bar = 5 μm. Enlarged views of the regions highlighted by yellow boxes are shown to the right of the main image. Scale bar = 300 nm. **(d)** Quantification of the nearest-neighbor distances between γH2AX and replication initiation signals from the experiments shown in (c). The control group shows the nearest-neighbor distances between γH2AX and EdU signals after randomizing the spatial distribution of γH2AX. **(e)** Representative images showing the distribution of γH2AX in WT (left) and KO (right) cells after 10 minutes of early S-phase. Scale bar = 5 μm. **(f)** Quantification of γH2AX fluorescence intensity in both cell types from the experiments shown in (e). n ≥ 30 cells. **(g)** Flow cytometry analysis showing the proportion of cells in S-phase for WT and KO cells, with 6 independent replicates. For panel (b), (d), (f) and (g), data represent mean ± SE. Significance levels are indicated as follows:* P<0.05; ** P<0.01; **** P<0.0001.

Importantly, the increased origin density under Lamin A/C deficiency correlated with elevated DNA damage. Immunofluorescence staining of γH2AX at 10 min after S-phase release was significantly higher under Lamin A/C deficiency (**Fig. 2e, f**), in line with the Western blotting data (**Fig. 2a, b**). Additionally, flow cytometry analysis revealed that the S phase was prolonged in Lamin A/C-deficient cells relative to wild-type cells (**Fig. 2g**), aligning with observations under Lamin A/C knockdown in other cell lines^62^. These results suggest that over-activation of replication origins in Lamin A/C-deficient cells increases replication initiation-induced DNA damage and genomic instability, extending the time required for genome replication.

### Lamin A/C Deficiency Affects Chromatin Structure in Replication Domains and Increases Chromatin Dynamics

Higher-order chromatin architecture plays a critical role in regulating nuclear processes, including DNA replication^18,32,33^. To explore whether Lamin A/C deficiency affects chromatin architecture, we first performed Hi-C sequencing to analyze chromatin interactions at the TAD scale in both wild-type and Lamin A/C-deficient cells. Under Lamin A/C deficiency, long-range chromatin interactions were reduced, while short-range interactions were increased (**Fig. 3a, b**). Furthermore, 5.6% of genomic regions in wild-type (WT) cells shifted from the A compartment to the B compartment in lamin A/C-deficient cells, whereas 8.5% of regions exhibited the reverse shift (**Supplementary Fig. 6a, b**). Notably, while TAD boundaries remain unchanged (**Supplementary Fig. 6c**), significant increase in chromatin interactions was detected at the level of TADs (**Fig. 3c**). Given the alignment between replication domains and TADs^28^, these results suggest that Lamin A/C deficiency enhances spatial chromatin interactions within replication domains.

**Figure 3.**
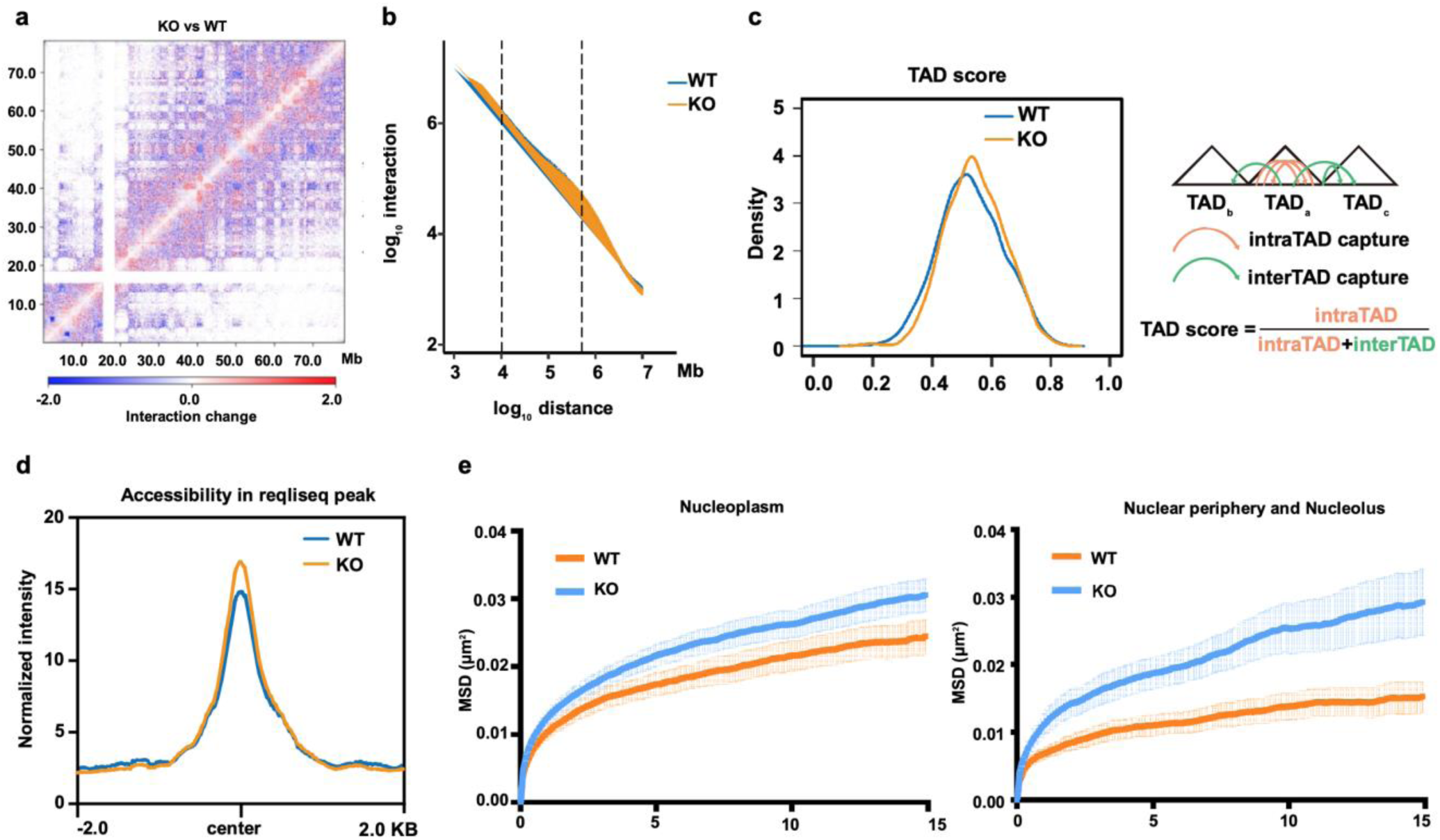
Regulation of Chromatin Dynamics and Structure by Lamin A/C Depletion. **(a)** Difference map of Hi-C interaction frequencies for chromosome 18 (in megabases) between WT (WT) and Lamin A/C knockout (KO) samples(MD-MBA-231), at a resolution of 100 Kb. The color gradient from blue to red represents the change in chromatin interactions, with blue indicating a decrease and red indicating an increase in interactions. **(b)** Hi-C interaction frequency as a function of increasing genomic distance (logarithmic scale) for WT and Lamin A/C-KO samples. Data are derived from two independent replicates. **(c)** Distribution of TAD scores in WT and Lamin A/C-KO samples. TAD scores are calculated as shown in the schematic: TAD score = intraTAD interactions / (intraTAD + interTAD interactions). Statistical significance (Wilcoxon test, p-value < 0.01, WT vs. KO). **(d)** Chromatin accessibility at BrdU-seq peaks for WT (blue) and Lamin A/C-KO (orange) samples (Hela S3), 2 independent experiments. **(e)** Averaged mean squared displacement (MSD) curves of chromatin loci dynamics in WT and Lamin A/C-KO cells for loci located in the nucleoplasm (left) and loci near the nuclear periphery and nucleolus (right) on chromosome 2 (n > 20).

Next, we performed ATAC-seq to assess changes in chromatin accessibility. The results indicated increased accessibility within the A compartment (**Supplementary Fig. 6d**), where early-replicating domains reside^26^. By integrating the ATAC-seq and BrdU-seq data, we observed a notable increase in chromatin accessibility at replicons in Lamin A/C-deficient cells (**Fig. 3d**), further supporting the idea that Lamin A/C plays a critical role in regulating replication domain structure.

A previous study reported that Lamin A/C regulates chromatin dynamics through tracking of telomeres^50^. To determine whether this regulation extends beyond telomeric regions, we performed single-molecule tracking of three CRISPR-SunTag-labeled genomic loci—one near the nuclear periphery, one in the nuclear interior, and one containing telomeres (Methods; **Supplementary Fig. 7a**). Under Lamin A/C-deficient conditions, all three loci exhibited increased mobility (**Fig. 3e; Supplementary Fig. 7b**). Extending previous studies that highlighted the relationship between chromatin dynamics and structural organization^63,64^, our findings demonstrated that the increased chromatin mobility in Lamin A/C-deficient cells was accompanied by significant increase of chromatin interactions and accessibility in replication domains, as revealed through Hi-C and ATAC-seq analyses.

Our previous research demonstrated that the spatial organization of RDs is closely linked to replication initiation efficiency^33^. To further investigate how Lamin A/C deficiency affects RDs’ spatial organization, we employed a similar metabolic labeling method for DNA replication (Methods), which labels the RDs for 45-min after the release from the G1/S phase. Using both STORM and super-resolution Expansion Microscopy (ExM; Methods), we imaged and analyzed the size of RDs. Notably, in Lamin A/C deficient cells, the sizes of RDs significantly increased (**Supplementary Fig. 8**). This finding aligns with observed increases in chromatin dynamics, intra-RD/TAD interactions, and chromatin accessibility, suggesting that Lamin A/C deficiency disrupts the structural integrity of RDs, thereby altering their spatial organization and functional properties.

### Lamin A/C Regulates Replication Initiation via Interacting with PCNA

Previous studies have revealed that PCNA interacts specifically with the Ig-fold domain at the C-terminus of Lamin A/C, and disrupting this interaction induces replication stress^57,58^. Accordingly, analyzing Lamin A/C interaction data from the BioGrid database^65^ also identified PCNA as a robust interacting partner of Lamin A/C (**Fig. 4a; Supplementary Fig. 9a**). To investigate how this interaction regulates DNA replication initiation, we first employed super-resolution microscopy to examine the spatial distribution of Lamin A/C and PCNA during the G1/S transition. Both PCNA and Lamin A/C within the nuclear interior were observed as punctate clusters (**Fig. 4b**). Interestingly, by calculating the nearest neighbor distance (NND) between PCNA and Lamin A/C, we identified two distinct populations of Lamin A/C clusters within the nuclear interior, contrasting with scrambled distributed Lamin A/C. A majority located 200 to 400 nm away from PCNA, exhibiting a distribution consistent with random chance, which suggests these clusters may serve PCNA-independent functions. However, a notable subset, representing 14.3% of the Lamin A/C clusters, was found in close proximity to PCNA, at distances of less than 100 nm (**Fig. 4b, c**). Expanding on the previous reports^58^, this proximity between Lamin A/C and PCNA suggests the regulation of their interaction in replication origin activation during the G1/S transition.

**Figure 4.**
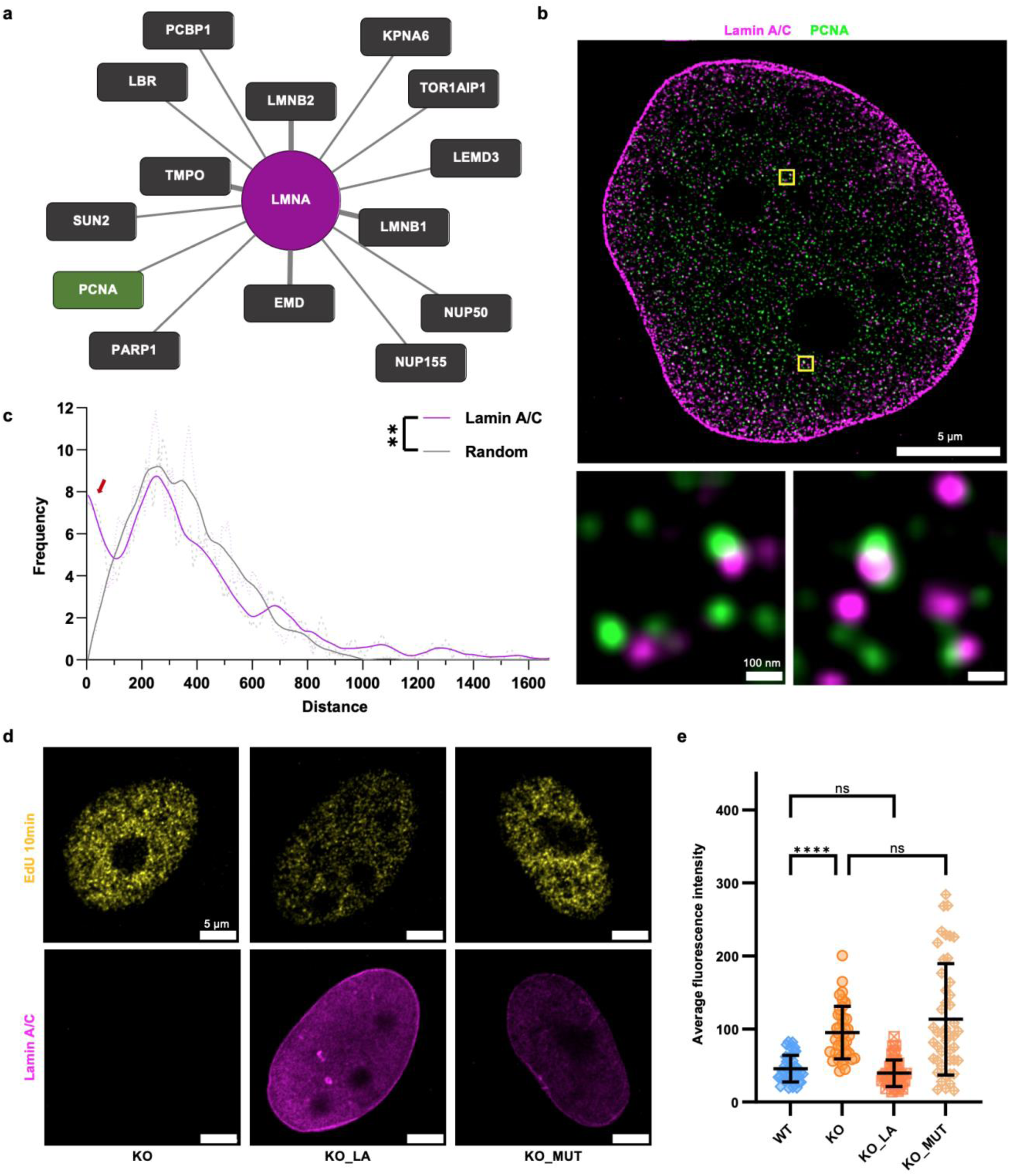
Interaction between Lamin A/C and PCNA regulates replication initiation. **(a)** Visualization of a selected interaction network of proteins identified in proximity to Lamin A/C(magenta), showing only proteins with an interaction evidence score greater than 8 (according to BioGRID and **Supplementary Fig. 9a**). PCNA is highlighted in green, as indicated in the figure key, to emphasize its role in the network. Other associated genes from the same organism(dark grey). thicker connection lines(grey) represent stronger evidence supporting the interactions. **(b)** STED microscopy image showing the spatial distribution of Lamin A/C (magenta) and PCNA (green) at the G1/S transition. The larger image on the top depicts the nuclear view. Scale bar = 5 μm. The lower panels show magnified views of the regions highlighted by yellow boxes in the larger image. Scale bar = 100 nm. **(c)** Frequency distribution of the nearest-neighbor distances between PCNA and Lamin A/C in non-perinuclear regions, based on the analysis in (b). The x-axis shows nearest-neighbor distances, and the y-axis shows frequency. The control represents randomized Lamin A/C distribution in non-perinuclear regions. Dashed lines show raw data, solid lines show lowess smoothed trends, and red arrows mark peaks of notable subset in the experimental group. Data are based on more than 20 cells. Significance levels are indicated as follows: ** P<0.01. **(d)** Representative images showing the distribution of replication initiation signals in Lamin A/C knockout cells (left), Lamin A/C WT rescue cells (middle), and Lamin A/C mutant rescue cells (right). After synchronization, cells were labeled with EdU for 10 minutes during early S-phase. Scale bar = 5 μm. **(e)** Quantification of the intensity of replication initiation signals per unit area in (d). Data are presented as mean ± SE, with n ≥ 30 cells. Significance levels are indicated as follows: ns, P>0.05; **** P<0.0001.

To further elucidate the role of the interaction between PCNA and Lamin A/C during replication initiation, we conducted rescue experiments in Lamin A/C knockout cells by expressing either wild-type Lamin A/C or Ig-fold deletion Lamin A/C, a mutant lacking the interaction with PCNA. Reintroducing wild-type Lamin A/C restored replication initiation activity to wild-type levels (**Fig. 4d, e**). In contrast, the reintroduction of the Lamin A/C Ig-fold deletion mutant resulted in replication initiation signals comparable to those observed in Lamin A/C knockout cells, with significantly higher levels than in wild-type cells (**Fig. 4d, e**). These findings underscore the essential role of the interaction between Lamin A/C and PCNA in regulating replication initiation and the maintenance of genome stability.

### Loss of Lamin A/C Increases PCNA Recruitment to Replication Domain

In our previous study, we showed that PCNA is maintained at the periphery of replication domains from G1 to G1/S, suggesting that its positioning is crucial for replication initiation^33^. Given this, we hypothesized that Lamin A/C deficiency might alter the spatial distribution of PCNA due to its interaction with Lamin A/C. To test this hypothesis, we followed a labeling and analysis protocol based on our previous work (Methods). Cells were first synchronized and released into the S phase, with EdU incorporated for 45 minutes to label early-replicating domains. In the subsequent cell cycle, the cells were re-arrested at the G1/S boundary and labeled for both EdU and PCNA. After imaging using expansion microscopy (ExM), the spatial distribution of PCNA relative to RDs was then analyzed using the Barycenter distance distribution (BDD) calculation.

Expanding on our prior findings^33^, we confirm that PCNA was primarily located at the periphery of RDs in wild-type cells during the G1/S transition (**Fig. 5a, b**), and this spatial organization remained largely unchanged in Lamin A/C-deficient cells, with BDD values both close to 1.

**Figure 5.**
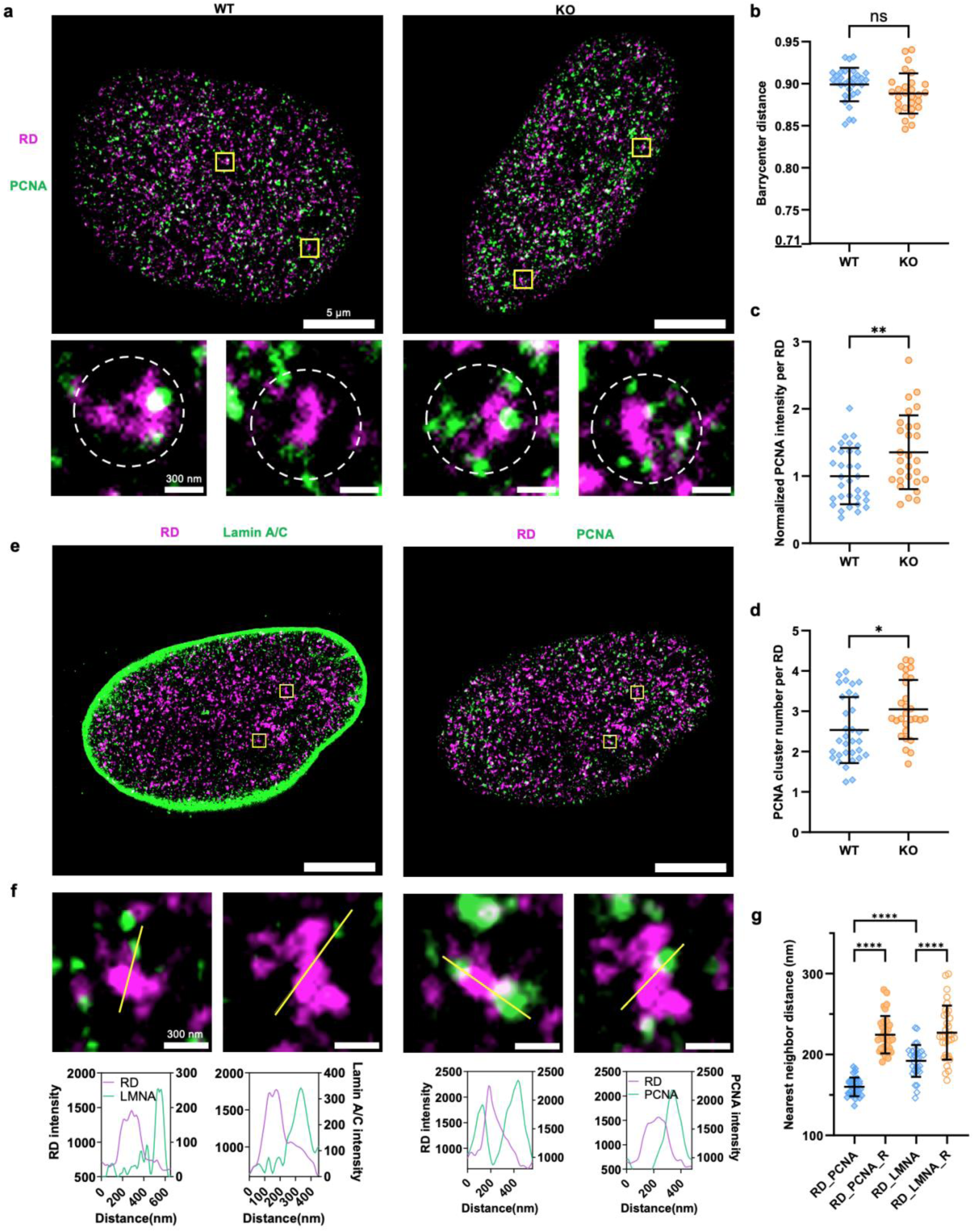
Spatial Distribution of Replication Domains (RD), PCNA, and Lamin A/C in WT and Lamin A/C Knockout Cells. **(a)** Representative Expansion Microscopy images showing the spatial distribution of PCNA(green) and replication domains (RD, magenta) in WT (left) and Lamin A/C knockout (right) cells. Scale bar = 5 μm. Enlarged rendered views of the regions highlighted by yellow boxes are shown to the below of the main image, with the white circle indicating the target RDs. Scale bar = 300 nm. **(b)** Dot plot quantifying the normalized barycenter distances between the centers of PCNA and RD in WT and Lamin A/C knockout cells from the experiments shown in (a). **The value 0.71 represents the BDD value by random simulations**^33^**. (c)** Dot plot quantifying the total PCNA fluorescence intensity near the RD (within 1.5 times the radius of gyration of the RD) in WT and Lamin A/C knockout cells from the experiments shown in (a). **(d)** Dot plot showing the number of PCNA clusters near the RD (within 1.5 times the radius of gyration of the RD) in WT and Lamin A/C knockout cells from the experiments shown in (a). **(e)** Representative Expansion Microscopy images showing the spatial colocalization of replication domains (RD, magenta) with Lamin A/C (green, left) or PCNA (green, right) in WT cells. Scale bar = 5 μm. **(f)** Enlarged rendered views of the yellow-boxed regions in (e). The left panels show the colocalization of RD (magenta) with Lamin A/C (green), and the right panels show the colocalization of RD (magenta) with PCNA (green). The line graphs below each image show the fluorescence intensity profiles along the yellow line in the corresponding image above. **(g)** Dot plot quantifying the nearest-neighbor distances between the centroids of RD and PCNA or Lamin A/C clusters from the experiments shown in (e). The random group represents the nearest-neighbor distances between RD and randomly distributed PCNA or Lamin A/C clusters. For panels (b), (c), (d) and (g), data are presented as mean ± SE, with n ≥ 30 cells. Significance levels are indicated as follows: ns, P>0.05; * P<0.05; ** P<0.01; **** P<0.0001

Notably, further analysis revealed that Lamin A/C deficiency influenced the enrichment of PCNA near replication domains. Specifically, we observed an increase in the number of PCNA clusters and an overall increase in PCNA fluorescence intensity near replication domains under Lamin A/C-deficient conditions (**Fig. 5c, d**). This finding supports the observed increase in replication origins (**Fig. 1**).

Additionally, to further validate these observations, CUT&Tag^66^ (Methods) experiments demonstrated a significant enhancement of PCNA enrichment to chromatin in Lamin A/C-deficient cells (**Supplementary Fig. 10a**). These results suggest that Lamin A/C deficiency promotes replication initiation by increasing PCNA recruitment to replication origins. All findings regarding the spatial distribution of replication domains and PCNA were independently validated using STED microscopy, with consistent results observed across replicates (**Supplementary Fig. 11**). Interestingly, when combined with the observed reorganization of RD structure (**Fig. 3**), the upregulation of PCNA enrichment suggests that the increased chromatin accessibility of RDs to the replication machinery facilitates increased recruitment of replication components at RD.

To further examine the relationship between RDs, PCNA, and Lamin A/C, we conducted simultaneous imaging of all three components at the G1/S transition. This analysis revealed that both Lamin A/C and PCNA were in close spatial proximity to RDs (**Fig. 5e-g**). On one hand, these results suggest that Lamin A/C plays a regulatory role in shaping RD structure, further supporting our previous findings (**Fig. 3**) by providing additional evidence on spatial distribution. On the other hand, PCNA was positioned ∼40 nm closer to RDs than Lamin A/C. This suggests that PCNA’s positioning at RD boundaries is critical for replication initiation and may be influenced by RDs’ structural changes in Lamin A/C-deficient cells.

### Lamin A/C Modulates the Availability of PCNA to Replication Origins

Previous studies have showed that the Lap2α–Lamin A/C complex regulates PCNA expression by modulating pRb and E2F1 activity^67,68^, suggesting that Lamin A/C influences PCNA expression. Beyond controlling expression, Lamin A/C also acts as an immobile nuclear scaffold protein that directly interacts with PCNA. This interaction raises the possibility that it sequesters free PCNA molecules within the nucleus and regulates their availability. Given this role, Lamin A/C depletion is expected to not only upregulate total PCNA expression but also increase the proportion of free PCNA.

To investigate this hypothesis, we first examined PCNA expression by quantifying its levels at both RNA and protein levels, observing a significant elevation compared to wild-type conditions (**Fig. 6a-c**). Notably, we also observed elevated PCNA immuno-fluorescence intensity during the G1/S transition, coinciding with increased replication initiation signal (**Fig. 6d, e**). These findings provide additional evidence supporting the regulatory role of Lamin AC in PCNA expression. Furthermore, quantitative analysis revealed that PCNA expression was positively correlated with the replication initiation signal, linking PCNA upregulation to replication origin activation (**Fig. 6f**).

**Figure 6.**
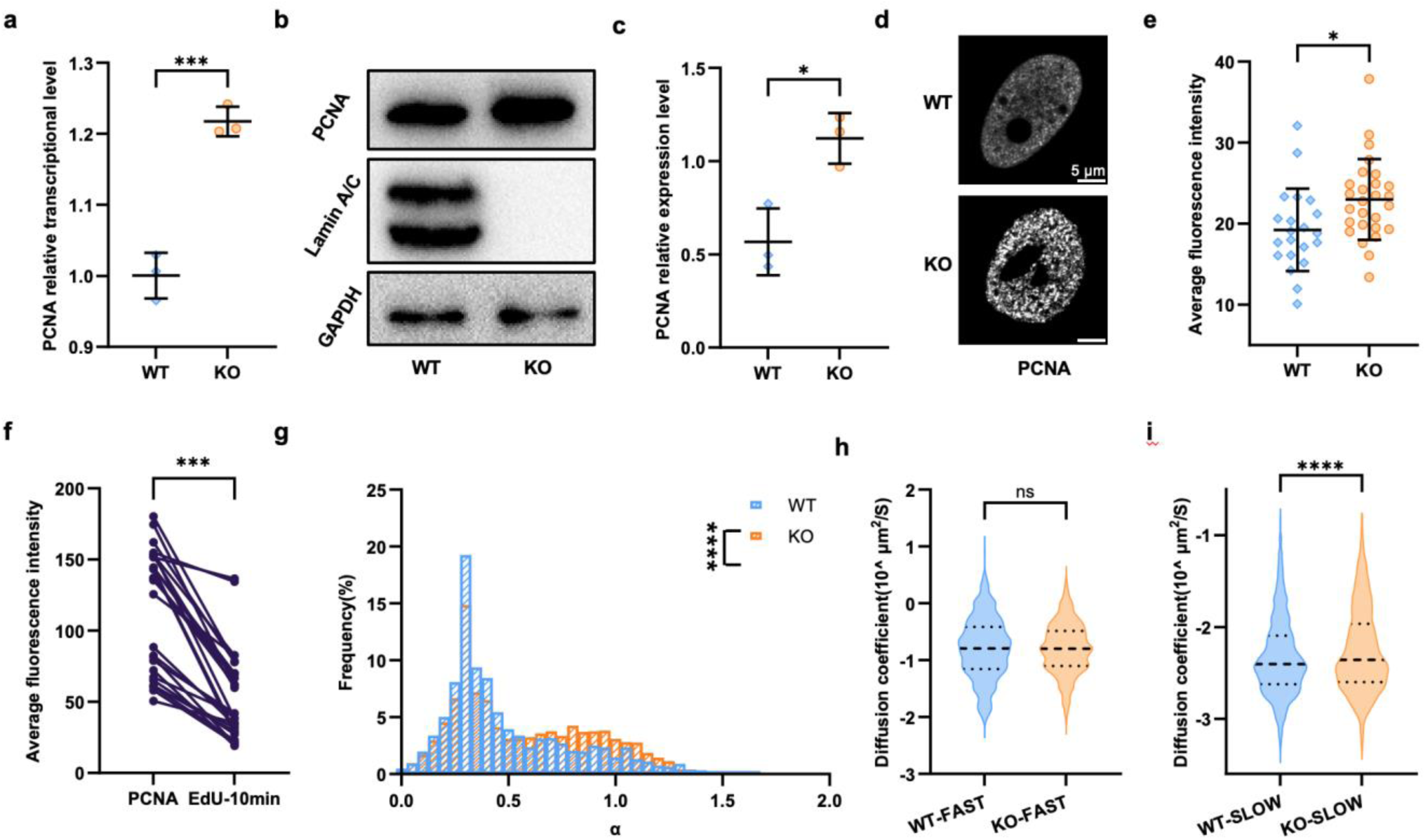
Impact of Lamin A/C depletion on PCNA dynamics and expression. **(a)** Quantification of relative PCNA transcriptional levels in WT and Lamin A/C knockout cells using reverse transcription quantitative PCR (RT-qPCR) (n = 3). **(b)** Representative immunoblot showing PCNA expression levels in WT and Lamin A/C knockout cells. **(c)** Quantification of relative PCNA protein levels from the experiments shown in (b) (n = 3). **(d)** Representative images showing the nuclear distribution of PCNA at the G1/S transition in WT (top) and Lamin A/C knockout (bottom) cells. Scale bar = 5 μm. **(e)** Quantification of nuclear PCNA fluorescence intensity per unit area from the experiments shown in (d), with n > 20 cells. **(f)** Scatter plot showing the correlation between the mean fluorescence intensity of PCNA and 10-min EdU labeling signals in WT cells. Each dot represents an individual cell. Statistical analysis is based on correlation analysis, with a R-value of 0.6657. *** P<0.001. **(g)** Frequency histogram plot of the anomalous diffusion index (α) for PCNA molecules during the G1/S transition in both cell types. **(h)** Violin plot showing the diffusion coefficients of DNA-bound PCNA molecules (α < 0.7) at the G1/S transition in both cell types. **(i)** Violin plot showing the diffusion coefficients of free PCNA molecules (α > 0.7) at the G1/S transition in both cell types. For panels (a), (c) and (e), data represent mean ± SE. For panels (h) and (i), data are shown as median ± quartiles. For panel (G-I), data are from more than 7 cells. For panels (a), (c), (e), (h) and (i), significance levels are indicated as follows: ns, P>0.05; * P<0.05; *** P<0.001; **** P<0.0001.

Next, to measure the availability of PCNA to replication origins, we generated stable cell lines expressing HaloTag-fused PCNA in both wild-type and Lamin A/C knockout cells to track the mobility of single PCNA molecules at the G1/S transition in real-time (**Supplementary Fig. 12**). HaloTag, a modified dehalogenase that binds to ligands conjugated with fluorescent dyes, enables precise live-cell protein tracking^69^.The analysis of PCNA’s anomalous diffusion coefficient (α value), reflecting molecular diffusion patterns, enabled us to categorize PCNA into two populations: a small proportion of freely diffusing molecules (α > 0.7) and a larger proportion of restricted molecules (α < 0.7), the latter group indicating PCNA molecules bound to DNA. Interestingly, in Lamin A/C-deficient cells, the proportion of freely diffusing PCNA increased significantly (**Fig. 6g**). Estimating DNA-bound PCNA revealed a ∼1.6-fold increase e in Lamin A/C-deficient cells compared to wild-type cells, in line with the enhanced PCNA enrichment towards RDs (Fig. 5c). These findings highlight that Lamin A/C deficiency increases the pool of freely diffusing PCNA proteins. Coupled with the elevated PCNA expression, this enhances PCNA’s availability and recruitment to replication origins.

Further analysis of the diffusion coefficient (D) for the distinct PCNA populations revealed that, the mobility of freely diffusing PCNA remained comparable between wild-type and Lamin A/C-deficient cells (**Fig. 6h**). Notably, the restricted PCNA population, presumably DNA-bound, exhibited increased mobility in Lamin A/C-deficient cells, reflecting a more dynamic chromatin environment (**Fig. 6i**). This enhanced mobility corresponds with the earlier observations of increased chromatin dynamics and structural changes upon Lamin A/C depletion (**Fig. 3**).

### Proposed Model for Lamin A/C Regulation of Replication Initiation and Genomic Stability

Based on our findings, we propose the following model to explain the role of Lamin A/C in regulating replication initiation and maintaining genomic stability (**Fig. 7**). Under normal conditions, Lamin A/C plays a dual role in ensuring the precise regulation of replication initiation. First, it restricts chromatin movement, stabilizing the spatial organization of replication domains and limiting the exposure of potential replication origins. Second, Lamin A/C regulates the availability of PCNA by interaction through the Ig-fold domain, sequestering a portion of PCNA and thereby reducing the number of freely available PCNA molecules. These regulations ensure that PCNA binds only to a limited number of replication origins, facilitating controlled replication fork progression and maintaining replication initiation balance.

**Figure 7.**
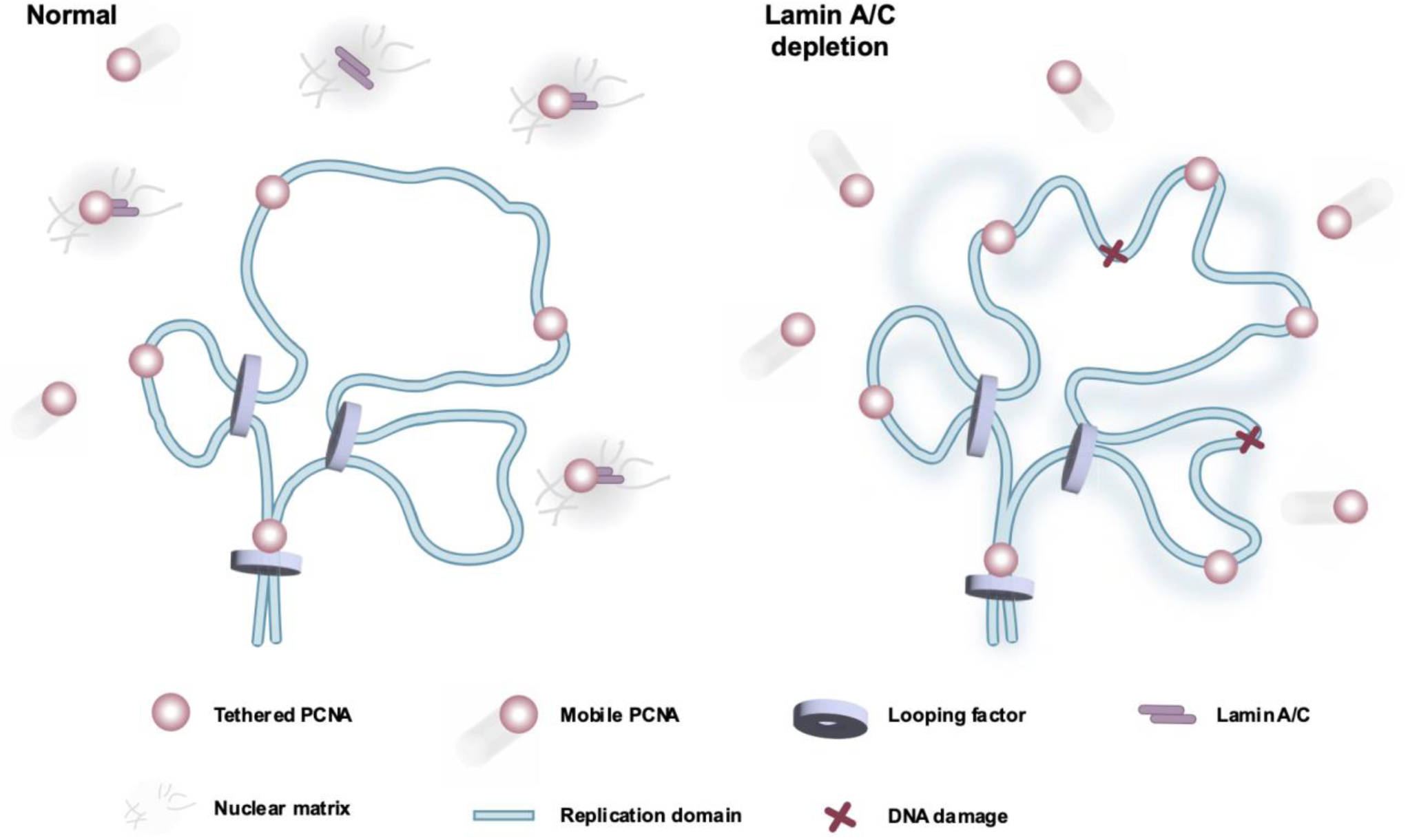
Lamin A/C regulates replication initiation and genomic stability. At the G1/S transition, the replication domain (blue) is surrounded by a nuclear scaffold containing Lamin A/C (purple sheets) and PCNA molecules (pink spheres), some of which are bound to replication domains, while others remain unbound. **Left (WT conditions):** Under normal conditions, replication domains are spatially stable, and unbound PCNA exists in two states: a freely diffusing pool (shaded pink spheres) and a portion sequestered by Lamin A/C on the nuclear scaffold (gray region). Only a limited number of PCNA molecules bind to activated replication origins, ensuring controlled replication initiation. **Right (Lamin A/C knockout conditions):** In Lamin A/C knockout cells, replication domain exhibits increased dynamics (blue shadow) and structural reorganization. Meanwhile, PCNA levels are upregulated, and Lamin A/C-mediated sequestration is lost, resulting in more freely diffusing PCNA. The disrupted replication domain structure and elevated free PCNA both contribute to an increased number of replication origins, leading to excessive replication initiation. This imbalance induces replication stress and genomic instability, resulting in DNA damage (red marks) in proximity to origins.

In the absence of Lamin A/C, two critical changes occur that disrupt this balance. First, the increased chromatin dynamics and spatial reorganization of RDs expose more replication origins to replication machinery. Second, the elevated levels of PCNA, coupled with the lack of Lamin A/C-mediated sequestration, lead to a higher proportion of freely diffusing PCNA. These freely available PCNA molecules are more likely to bind to the newly exposed replication origins, resulting in an increase in the number of active replication initiation events. This imbalance in replication origin activation subsequently leads to replication stress, DNA damage, and genomic instability.

### Discussion and Perspective

Our study demonstrates that Lamin A/C deficiency significantly impacts DNA replication initiation and chromatin structure. Firstly, Lamin A/C deficiency led to an increased number of active replication origins (**Fig. 1**) in the early S-phase. Consequently, these changes in replication initiation contributed to increased DNA damage, as evidenced by elevated γH2AX levels, leading to genomic instability (**Fig. 2**). The increased replication initiation was accompanied by accelerated chromatin dynamics, altered chromatin structure, resulting in enhanced accessibility near replication origins (**Fig. 3**). Moreover, as a key component of the replication machinery, PCNA was identified to interact with Lamin A/C, via its Ig-fold domain mediating this interaction and playing a crucial role in regulating DNA replication initiation (**Fig. 4**). Super-resolution imaging revealed that Lamin A/C deficiency results in enlarged replication domains, disrupting their spatial organization and enhancing accessibility, which likely facilitated greater PCNA recruitment (**Fig. 5**). Lamin A/C deficiency also led to increased PCNA availability, promoting greater recruitment of PCNA to chromatin and replication domains (**Fig. 6**). Taken together, these findings support our proposed model (**Fig. 7**) in which Lamin A/C maintains genomic stability by simultaneously regulating the spatial organization of replication domains and the availability of replication factors, thus ensuring balanced replication initiation and preventing excessive replication origin firing. Previous studies highlighted Lamin A/C’s role in chromatin dynamics and nuclear architecture^50,51,53^, but its influence on higher-order chromatin structures, such as TADs/RDs, remains less understood. Our findings address this knowledge gap, showing that Lamin A/C, under normal conditions, restricts chromatin mobility and stabilizes RD structures to maintain their spatial organization. By preserving RD integrity, Lamin A/C ensures proper spatial regulation of replication origin activation. This supports our earlier work suggesting that RDs structure influences the likelihood of origin firing, with origins at RDs/TADs peripheries being more likely to activate^33^.

Our study investigates how Lamin A/C regulates replication initiation and maintains genomic stability by modulating chromatin accessibility and its interactions with replication machinery. These insights have broader implications for understanding Lamin A/C’s involvement in cancer and viral infections, including HPV. Additionally, Lamin A/C’s regulation of RD structure extends to its impact on key replication machinery components, specifically PCNA. Under normal conditions, Lamin A/C modulates the availability of PCNA by maintaining a controlled pool of free PCNA within the nucleus, thus fine-tuning its recruitment to RDs (**Fig. 5, 6**). This precise regulation is crucial for preventing excessive origin activation, thereby reducing the likelihood of replication stress. In contrast, Lamin A/C’s binding partner Lap2α, which is known to regulate PCNA expression levels^67,68^, exerts a much weaker and opposite effect on chromatin dynamics compared to Lamin A/C^70^. This distinction underscores the unique role of Lamin A/C in coordinating chromatin architecture and replication machinery to ensure spatial and functional control of DNA replication initiation.

Our study establishes a novel link between Lamin A/C regulation, replication origin selection, and replication-associated DNA damage. While Lamin A/C’s role in maintaining replication fork stability and facilitating DNA damage repair has been well-documented^55,71,72^, our findings highlight its direct influence on replication initiation, positioning it as a gatekeeper that restricts replication origin activation to mitigate replication stress and maintain genomic stability. Notably, replication origin activation is tightly linked to replication timing, which is disrupted by ATR/Chk1 dysfunction, leading to premature origin activation within mid-to-late RDs^73,74^. In contrast, Lamin A/C deficiency increases the number of active replication origins without altering replication timing, ultimately resulting in elevated DNA damage. (**Fig. 1, 2**). Furthermore, the clustering of upregulated origins near TAD boundaries (**Supplementary Fig. 3d, e**) implies that Lamin A/C may preferentially target conserved origins, rather than randomly activating flexible origins within RDs, though further studies are needed to confirm this specificity. Beyond its role in replication, Lamin A/C deficiency or dysfunction also results in nuclear softening, causing DNA damage under mechanical stress^75,76^. This mechanical vulnerability likely explains the moderate increase in DNA damage observed in the G1 phase (Fig. 2a, b), where replication-associated stress is absent. Together with excessive replication origin activation during the early S phase, these complementary mechanisms contribute to genome instability in Lamin A/C-deficient cells.

The link between Lamin A/C dysregulation and DNA damage suggests broader implications, particularly for cancer and viral infections. As genomic instability is a hallmark of cancer, the increased DNA damage observed in Lamin A/C-deficient cells offers a potential mechanistic connection to tumorigenesis. Additionally, the prolongation of the S phase in Lamin A/C-deficient cells creates a replication environment conducive to viral exploitation. For example, HPV, which is known to downregulate Lamin A/C expression^56^, relies on the host DNA replication machinery for its replication cycle. The prolonged S phase observed in Lamin A/C-deficient cells may facilitate viral replication and promote the integration of the viral genome into the host genome, thereby linking Lamin A/C loss to both genomic instability and heightened susceptibility to HPV infection. Together, these findings underscore the critical role of Lamin A/C in maintaining genomic integrity and protecting against both cancer development and viral exploitation.

While our study provides valuable insights into Lamin A/C’s regulation of replication initiation and genome stability, several questions remain unanswered. Beyond its impact on RD structure and PCNA availability, it is unclear whether Lamin A/C also influences other key replication factors, such as ORC and MCM. Understanding its broader role in origin licensing could uncover new layers of replication control. Additionally, although we observed increased PCNA availability, it remains unclear whether Lamin A/C also modulates PCNA loaders/unloaders^10,77,78^ to fine-tune PCNA recruitment at the replication sites. Clarifying this mechanism could further explain its role in maintaining replication fidelity.

Moreover, despite identifying the Ig-fold domain of Lamin A/C as critical for its interaction with PCNA, mapping the precise binding interface on PCNA itself is still needed. Advanced protein interaction mapping and structural analyses could address this gap. Finally, linking our findings to *in vivo* models would provide insights into Lamin A/C’s role in disease contexts, such as cancer and viral infections. For instance, HPV, which downregulates Lamin A/C, might exploit the replication environment to enhance viral genome integration, highlighting the therapeutic potential of targeting Lamin A/C pathways in replication-associated diseases.

## Materials and Methods

### Cell culture

HeLa S3 cells and MDA-MB-231 cells were cultured in 10 cm plastic dishes (Corning, 430167) or 35 mm glass-bottom dishes (Cellvis, D35-20-1.5-N) to facilitate monitoring and imaging. The cells were maintained in Dulbecco’s Modified Eagle Medium (DMEM, Gibco, 11965-092) supplemented with 10% fetal bovine serum (Gibco, 10099-141) and 100 U/mL penicillin-streptomycin (Gibco, 15140-122). Cultures were incubated at 37°C in a humidified atmosphere containing 5% CO₂ (Thermo Scientific, Forma Series II). Cell growth was monitored regularly, and subculturing was performed every 2-3 days when confluency reached approximately 80%, using 0.25% trypsin (Gibco, 25200-056).

### Cell synchronization

To synchronize the cell cycle at the G1/S transition, a combination of thymidine and aphidicolin was employed (**Supplementary Fig. 2a**). Cells were treated with 2 mM thymidine (Sigma-Aldrich, T1895) for 16 hours to block DNA synthesis. Following three washes with PBS (Gibco, C14190500BT), cells were incubated in fresh DMEM for 10 hours. Aphidicolin (Sigma-Aldrich, A0781) was then added at a final concentration of 2 μg/mL for 16 hours, effectively enhancing the synchronization at the G1/S boundary. The following release was conducted using 10 μM EdU or 16 μM BrdU in DMEM, with varying incubation duration, as described respectively in the article.

In the non-aphidicolin-induced approach (**Supplementary Fig. 4a**), cells were synchronized in the M phase using nocodazole (Sigma-Aldrich, M1404), which inhibits spindle formation. Cells were incubated with 100 ng/mL nocodazole for 16 hours. Detached mitotic cells were collected by gently shaking the culture dishes, and after centrifugation, the cells were washed three times with PBS. The cells were subsequently released into DMEM containing 10 μM EdU (Thermo Fisher Scientific, C10337) for 8 hours to label cells entering early S phase.

### Click chemistry and immunofluorescence

Click chemistry and immunofluorescence staining was used to detect DNA replication events and specific proteins. Cells were fixed and washed with PBS. To reduce non-specific binding, the samples were blocked in PBS containing 5% bovine serum albumin (BSA, Sigma-Aldrich, A7906) and 0.5% Triton X-100 (Sigma-Aldrich, T8787) for 45 minutes.

For click chemistry, the Click-iT EdU Imaging Kit (Thermo Fisher Scientific, C10337) was used to label EdU with the included AF647 dye for STORM experiments or with 10 μM SIR dye for other experiments. When SIR dye was used for clicking, DMSO was added at a final concentration of 10% to enhance dye isolation.

For immunofluorescence, Primary antibodies against BrdU (BD, B44), PCNA (CST, 13110), Lamin A/C (CST, 4777), pH2AX (CST, 9718) and CTCF (Abcam, ab128873) were diluted 1:200 in PBS with 5% BSA and 0.5% Triton X-100. The samples were incubated with primary antibodies at room temperature for 1 hour, followed by four washes with PBS containing 1% Triton X-100. Secondary antibodies conjugated to different dyes, chosen to suit specific imaging experiments (STED, ExM and STORM), were applied at a 1:50 dilution and incubated for 1 hour. After another series of washes, the samples were stored in PBS at 4°C, protected from light until imaging.

### DNA fiber assay

The DNA fiber assay was used to measure replication fork progression in synchronized cells. Hela cells were synchronized at the G1/S transition and released into S phase by washing with PBS three times, followed by incubation in DMEM containing 40 μM CIdU and then 100 μM IdU each for 20 minutes. Cells were harvested using trypsin, resuspended in PBS, and collected by centrifugation. The cell pellet was washed three times in PBS, and 100 μL of lysis buffer (200 mM Tris-HCl (Sigma-Aldrich, T5941), 50 mM EDTA (Sigma-Aldrich, E6758), and 0.5% SDS (Sigma-Aldrich, L3771)) was added.

For DNA combing, lysed DNA was applied to silanized coverslips and allowed to dry at room temperature for 3-5 minutes. Coverslips were tilted at an approximately 15-degree angle to allow gradual flow of the buffer, ensuring uniform DNA stretching. After air drying, the DNA was fixed using a 3:1 methanol (Sigma-Aldrich, M1775)/acetic acid (Sigma-Aldrich, A6283) mixture. Newly synthesized DNA was labeled using immunofluorescence with anti-CIdU (BD, B1/75) and anti-IdU (BD, B44) antibodies. Fluorescence microscopy was used to image the labeled DNA fibers, and the lengths were measured manually. Replication fork speed was calculated by converting physical fiber lengths into kilobases using a factor of 2 Kbp/μm.

### Flow cytometry analysis

Flow cytometry was performed to quantify cell cycle progression. Hela cells were incubated in 30 μM BrdU (Sigma-Aldrich, B5002) for 60 minutes, then washed with PBS and trypsinized. After resuspension in DMEM containing 10% fetal bovine serum (FBS; Sigma-Aldrich, F2442), cells were collected by centrifugation at 200 g for 3 minutes. They were washed twice with PBS and fixed by adding ice-cold 70% ethanol in a 3:1 ratio with PBS, followed by storage at -20°C overnight.

The next day, cells were washed twice with PBS, resuspended in PBS, and treated with 100 μg/mL RNase A (Sigma-Aldrich, R6513) and 0.2% Triton X-100 (Sigma-Aldrich, X100) at 37°C for 30 minutes. Propidium iodide (PI; Solarbio, P8080-10) was added at a final concentration of 50 μg/mL, and the cells were incubated for 30 minutes at room temperature. Flow cytometry was conducted using a BD analyzer, with a minimum of 10,000 cells collected per sample. Debris and doublets were excluded by gating on FSC-A/SSC-A and PE-A/PE-W plots. Data were analyzed to determine the proportion of cells in the S phase, with voltage adjustments made to ensure accurate separation of G1, S, and G2/M phases in PI-A histograms.

### BrdU-Seq

BrdU-seq (5-bromo-2’-deoxyuridine sequencing) was modified from the previous study^79^ for replication origin detection, cells were synchronized at the G1/S boundary and labeled for 10 minutes to correspond with imaging results.

Hela cells were cultured in 10 cm dishes at a density of ∼20%, then synchronized at either the G1/S boundary or specific points in S phase. After synchronization, cells were washed three times with PBS. BrdU (Sigma-Aldrich, B5002) was added to the medium to label replicating DNA for specific durations. Following labeling, cells were washed with PBS, and lysed in RIPA buffer (Yeasen, 20101ES60) containing RNase A (Qiagen, 19101) and Proteinase K (Thermo Fisher Scientific, EO0491) at final concentrations of 0.1 mg/mL and 0.2 mg/mL, respectively. Lysis was carried out in a thermomixer first at 37°C for 2 hours, followed by 55°C for 5 hours to ensure complete cell lysis and DNA release.

Genomic DNA was extracted from the lysate using phenol-chloroform extraction to ensure purity. DNA was mixed with phenol-chloroform-isoamyl alcohol (24:25:1) at a 1:1 ratio, shaken for 5 minutes, and allowed to settle for phase separation. After centrifugation (15,000 g, 10 minutes), the upper aqueous phase containing the DNA was collected. The extraction was repeated, and the DNA was precipitated with ethanol and 3 M sodium acetate (pH 5.5). DNA was pelleted by centrifugation (16,000 g, 30 minutes at 4°C), washed with 75% ethanol, air-dried, and resuspended in ultrapure water.

The extracted DNA was fragmented into ∼200 bp fragments using a Covaris sonicator (175 W, 10% duty cycle, 200 cycles per burst). Fragmented DNA was then purified using Vazyme magnetic beads (Vazyme, N401-01). Following DNA fragmentation, BrdU-labeled DNA was denatured to single-stranded DNA by heat treatment. The denatured DNA was incubated with BrdU-specific antibodies (BD, 555627) in an immunoprecipitation buffer. Protein G-Dynabeads (Invitrogen, 10004D) were used to capture the antibody-DNA complexes. The immunoprecipitated DNA was washed extensively, and DNA was eluted with TE buffer. The recovered DNA fragments were used to generate sequencing libraries with the Vazyme single-strand DNA library preparation kit (Vazyme, NE103). Libraries were sequenced on an Illumina platform.

Sequencing data were preprocessed to remove low-quality reads, and adapter sequences were trimmed using Trim Galore. Processed reads were aligned to the human reference genome (hg19) using Bowtie2. Peak calling was performed using MACS2^80^ to identify BrdU-enriched regions. Visualization of BrdU-labeled regions was done using the Integrative Genomics Viewer (IGV). Differential analysis between wild-type and Lamin A/C knockout cell lines was conducted using DiffBind and DESeq2^81^, with significant peaks identified by thresholds of p < 0.05 and log2FoldChange > 1 or < -1. When considering the reported fork speed (∼1 Kbp /min)^82,83^, an estimated 10-min replication peak length of ∼10 Kbp was chosen to select the peak containing at least one replication origin.

### Hi-C

Hi-C was performed as described previously^23^. Hela cells were cross-linked with 3% formaldehyde at room temperature for 10 minutes, followed by quenching with 0.125 M glycine. Nuclei were isolated, and chromatin was digested using a restriction enzyme, HindIII, for 16 hours at 37°C. Digested DNA ends were biotinylated and ligated in situ to create proximity ligation products. Cross-links were reversed, and the DNA was purified. Biotin-labeled ligation junctions were isolated using streptavidin-coated beads, and the libraries were prepared using standard Illumina sequencing protocols.

Sequencing reads were aligned to the reference genome hg19, and interaction maps were generated using computational tools HiC-Pro^84^. Hi-C raw matrices were normalized by ICE (Imakaev et al., 2012). After normalization, the genome-wide relative contact maps are comparable among each sample, the subsequent analysis were based on the normalized matrix. The genome was partitioned into A/B compartments according to the first principal component values: positive values defined compartment A (higher gene density), while negative values defined compartment B. For TAD analysis, we utilized ICE-normalized matrices at a 40-kb resolution to identify TADs using the Perl script matrix2insulation.pl (http://github.com/blajoie/crane-nature-2015).

### ATAC-seq

ATAC-seq was performed to assess chromatin accessibility as previously described^85^. Briefly, nuclei were extracted from G1/S-transition synchronized cells using lysis buffer and centrifuged at 500×g for 5 minutes at 4°C. Nuclei pellets were resuspended in the Tn5 transposase reaction mix and incubated at 37°C for 30 minutes. Following transposition, equimolar adapters were ligated, and the resulting libraries were amplified by PCR. Libraries were purified using AMPure beads and quantified using a Qubit fluorometer. Sequencing was conducted on an Illumina NovaSeq 6000 platform with 150 bp paired-end reads.

Raw sequencing reads were processed to remove adapters, poly-N sequences, and low-quality reads (base quality <15 for >40% of reads). Clean reads were aligned to the reference genome hg19 using Bowtie2. Reads derived from mitochondrial DNA or with low mapping quality (MAPQ <13) were excluded. Only uniquely mapped and de-duplicated reads were retained for analysis. Peak calling was performed using MACS2 (v2.2.7.1) with the default parameters. Identified peaks were annotated using ChIPseeker to associate them with nearby genes, followed by Gene Ontology (GO) and KEGG pathway enrichment analyses to identify functional enrichment. Differential peak analysis between groups was performed using Bedtools^86^, with differential peaks defined as having a fold change > 2 (|log2FC| > 1). All experiments were conducted in triplicate to ensure reproducibility, and statistical significance was determined with a threshold of p < 0.05.

### Cut&Tag

The Cut&Tag protocol was adapted from previous study^66^ for efficient profiling of chromatin-bound factors. Cut&Tag was performed using the Vazyme CUT&Tag kit (#TD904) according to the manufacturer’s instructions, with minor modifications. Briefly, approximately 100,000 Hela cells were harvested and permeabilized with digitonin in wash buffer. ConA beads were activated and incubated with cells at room temperature for 10 minutes. Primary antibodies (PCNA) against the target protein were added, and the reaction was incubated overnight at 4°C. The next day, samples were washed and incubated with a secondary antibody conjugated to Protein A-Tn5 transposase. Following tagmentation, DNA was extracted using DNA Clean Beads (Vazyme #N411) and purified. The libraries were amplified using the Vazyme library amplification primers (#TD202/TD203) and sequenced on an Illumina NovaSeq 6000 platform. Sequencing reads were aligned to the reference genome, and peaks were called using MACS2^80^. The subsequent analysis process was the same as that of ATAC-seq.

### RNA-seq

Total RNA was extracted and its integrity assessed using the RNA Nano 6000 Assay Kit on the Bioanalyzer 2100 system (Agilent Technologies). mRNA was enriched using poly-T magnetic beads, fragmented, and reverse transcribed into cDNA. Following end repair, A-tailing, and adaptor ligation, 370–420 bp cDNA fragments were selected, PCR amplified, and subjected to quality control using a Qubit Fluorometer, Agilent 2100, and qRT-PCR to ensure effective library concentrations (>2 nM). Qualified libraries were sequenced on the Illumina NovaSeq 6000 platform, generating 150 bp paired-end reads. Raw data were processed to remove low-quality reads and adapters, producing high-quality clean reads. were mapped to the reference genome using STAR^87^, and gene expression levels were quantified as FPKM using featureCounts, accounting for sequencing depth and gene length. Differential expression analysis was performed using the DESeq2^81^ R package (v1.20.0) with two biological replicates per condition. Significant genes were identified based on padj < 0.05 and |log2(fold change) | > 1. Genes with p-value < 0.05 and fold change > 0.5 were further analyzed in DAVID (KEGG and GO analysis were focused). Terms with FDR < 0.05 were ranked by count, and the top 10 pathways with the highest counts were selected for display.

### Stable cell line construction

Lamin A/C Knockout cell line: sgRNAs targeting Lamin A/C (CGTCATGAGACCCGACTGG for Hela S3, CGTCATGAGACCCGACTGG and CATCGACCGTGTGCGCTCGC for MDA-MB-231) were respectively inserted into the lentiCRISPR v2 vector. L entiviral particles were generated by co-transfecting HEK293T cells with lentiCRISPR v2, psPAX2, and pMD2.G plasmids in a 5:3:2 ratio using PEI MAX 40000 (Polysciences, USA). The viral supernatants were harvested 48 h after transfection and concentrated by ultracentrifugation at 20,000 × g for 90 min at 4 °C. MDA-MB-231 cells were infected with the concentrated lentivirus in the presence of 8 μg/mL polybrene (Sigma-Aldrich, USA). Transduced cells underwent selection with 2 μg/mL puromycin 48 hours post-infection. Individual clones were isolated using flow cytometry, and the knockout of Lamin A/C was verified through Western blotting.

PCNA-HaloTag cell line: To study PCNA dynamics, a stable Hela cell line expressing PCNA-HaloTag was constructed. The sgRNA sequence, GACCAGGCGCGCCTCGAACA, was used to introduce a HaloTag at the PCNA locus via homology-directed repair. Hela cells were co-transfected with Cas9-sgRNA plasmids (adapted from Addgene-PX330) and a donor plasmid containing the HaloTag-PCNA homology arms (adapted from Addgene-170818). Following transfection, cells were labeled with JF594 HaloTag ligand and sorted using flow cytometry. Successful integration was confirmed by PCR and immunoblotting.

sfGFP Cell line for chromatin loci tracking: The construction was performed using plasmids and protocols identical to those described by our lab previous work^88^. Briefly, WT and Lamin A/C knockout Hela cells were seeded into 6-well plates at approximately 50% confluency one day prior to transfection. Each well was transfected with pB-TRE3G-NLSSV40-dCas9-3X NLSSV40-24X GCN4-V4-NLSSV40-P2A-BFP-PuroR-P2A-rtTA plasmid, pB-TRE3G-ScFV-sfGFP-GB1-NLSSV40-HygroR-P2A-rtTA plasmid, and pCAG-hyPBase plasmid. After 48 hours, dual antibiotic selection with 200 μg/mL hygromycin and 5 μg/mL puromycin was applied for two weeks. Single-cell clones expressing BFP and GFP were sorted by flow cytometry and expanded for subsequent chromatin dynamic tracking experiments.

### Single-Molecule Tracking (SMT) and data analysis of PCNA

To enable PCNA single-molecule tracking, target proteins were expressed as HaloTag fusion proteins in stable cell lines generated as described earlier. Hela cells were synchronized at the G1/S transition and labeled with the fluorescent dye JF549 (10 μM, diluted 1:1000 in DMEM containing aphidicolin at 2 μg/mL, Sigma-Aldrich, A0781). The cells were incubated at 37 °C for 20 minutes to allow the dye to bind to the HaloTag fusion proteins. After labeling, unbound dye was removed by washing the cells three times with PBS containing aphidicolin. The cells were then transferred to phenol red-free DMEM (also containing aphidicolin) for imaging. The presence of aphidicolin ensured that cells remained in the G1/S transition during the labeling and imaging processes. Single-molecule tracking was performed similarly to previously described^89^, using Nikon NSTORM microscope equipped with a 561 nm laser (2RU-VFL-P-300-488-B1R; MPB) and total internal reflection fluorescence (TIRF) capability. The microscope was adjusted to high-inclined laminated optical sheet (HILO) illumination to reduce nuclear background and enhance signal quality. Initial widefield fluorescence imaging was conducted to locate cells in the appropriate phase. After selecting appropriate cells, 100 frames of widefield images were captured with split lase intensity set as 1%. For single-molecule tracking, the split laser intensity was increased to 100% (laser power∼35 mW), with an exposure time of 20 ms. Rapid photobleaching was performed for approximately 30 seconds to reduce background signals and sparsely label individual molecules suitable for tracking. Once the background was sufficiently reduced and the single-molecule signal was sparse, 10,000-20,000 frames of single-molecule images were captured.

The acquired single-molecule trajectories were analyzed using Fiji with the TrackMate^90^ plugin. Single molecules were automatically detected and tracked using the Laplacian of Gaussian (LoG) detector. Parameters were adjusted to accurately detect molecules while excluding noise. TrackMate was used to process the image sequences and output single-molecule trajectories. The resulting trajectory data were exported to custom MATLAB scripts for further analysis. Each individual MSD curve was fitted by least-squares regression to the following model.

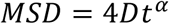

Diffusion coefficients (d) and anomalous diffusion exponents (α) were calculated to quantify the molecules’ movement. The diffusion coefficient was estimated by measuring the mean squared displacement (MSD) of molecules over time, representing their mobility. The anomalous diffusion exponent was used to assess deviations from Brownian motion.

### Chromatin loci dynamic tracking

Chromatin loci dynamic tracking was adapted from previously described^88^. WT and Lamin A/C KO Hela cells were transfected with sgRNA expression plasmids described above targeting the same genomic loci as previous work^88^. After 24–48 hours of culture, cells were prepared for imaging.

Dynamic tracking experiments were conducted on a Nikon N-STORM microscope with a 488 nm laser (2RU-VFL-P-300-488-B1R; MPB) to excite the sfGFP fluorophore. For chromatin loci dynamic tracking, the laser power was set to 10 µW, and images were acquired with 100 ms exposure time. Cells were maintained at 37 °C and 5% CO₂ in a stage-top incubation chamber to ensure viability during live-cell imaging. The further analysis was consistent with PCNA single molecular tracking.

### Confocal and STED microscopy imaging and data analysis

Confocal and STED imaging was performed using the Stedycon system (Abberior) equipped with a 100× oil-immersion objective (UPlanSApo 100×, 1.40 NA, Olympus) for high-resolution imaging. HeLa cells were seeded onto glass-bottom dishes and synchronized as described in the article. Immunofluorescence staining was performed, with or without prior EdU labeling, depending on the description in the Main. EdU labeling, where applicable, was achieved via click chemistry using the SIR dye. Specifically, STAR Orange (Abberior) and STAR Red (Abberior) dyes were used for STED immunofluorescence staining, corresponding to the green and magenta markers in the figure, respectively. Prior to imaging, samples were washed three times with ultrapure water and air-dried overnight, followed by mounting with an appropriate mounting medium. The mounted samples were stored at -20°C until imaging.

During imaging, the samples were positioned on the stage, and the focus and target cells were carefully adjusted to ensure optimal clarity in all channels. For STED, a 775 nm depletion laser was used in combination with either a 647 nm or 561 nm excitation laser, depending on the fluorophore, to capture confocal images. Key imaging parameters were optimized, and the system’s software was adjusted to achieve the best resolution and signal-to-noise ratio. The specific imaging parameters used in this experiment are detailed as follows: Replication origins (EdU, 10 min) and replication domains (EdU, 45 min) were visualized in the 647 nm channel with a STED laser intensity of 30% and a scan number of 5. Lamin A/C was also imaged in the 647 nm channel with a STED laser intensity of 40% and a scan number of 5. PCNA, CTCF, and pH2AX were imaged in the 561 nm channel with STED laser intensities of 60%, 60%, and 70%, respectively, and each with a scan number of 5. STED imaging was carried out with appropriate adjustments to the excitation and depletion laser intensities to minimize photobleaching while maintaining high-resolution signal capture.

STED images were deconvolved using Huygens Professional software (Scientific Volume Imaging, https://svi.nl/Huygens-Professional). System-adaptive parameters provided by the software were used to ensure accurate deconvolution while preserving the authenticity of the raw data. Deconvolved images were converted into TIFF format. The TIFF images (including confocal and STED) were analyzed using custom-developed code to identify and quantify specific biological structures. The analysis pipeline first involved the identification of nuclei using morphological algorithms that delineate the core regions of cells. Adaptive thresholding was then applied to segment protein clusters and replication signals. Several key parameters were computed to assess the features and spatial distribution of protein clusters within the cells: mean fluorescence intensity, cluster count, size, area, nearest neighbor distance (NND) and barycenter distance. The barycenter distance was defined as the spatial barycenter distance between the origins and the RD, normalized by the RD’s radius. To ensure accurate assignment, an origin was analyzed only if its spatial barycenter distance to the nearest RD was shorter than half of the RD’s major axis length.

### Expansion Microscopy (ExM) Imaging and Data Analysis

HeLa cells were seeded onto coverslips and synchronized, followed by 45-min EdU incorporation, click chemistry with dye SIR, and immunofluorescence for PCNA with dye AF594, as previously described. The ExM protocol was then adapted from the methods outlined by Shi et al^35^.

Following the sample preparation, cells were post-fixed in a solution containing 3% PFA and 0.1% glutaraldehyde (GA) in PBS for 15 minutes at room temperature with gentle shaking. This fixation buffer was freshly prepared by dissolving 4 g of PFA in 50 mL of water, heated to 60°C to assist dissolution, and adjusted to a pH of 7.0–7.4.

The cells were washed three times in PBS (5 minutes per wash), followed by a second fixation step using 0.7% PFA and 1% acrylamide (AaM) in 1× PBS at 37°C for 1–6 hours. This step ensured direct anchoring of proteins and DNA into the first layer of hydrogel. Cells were then incubated in a pre-gel solution composed of 19% sodium acrylate (SA), 10% acrylamide (AAm), 0.1% N,N’-methylenebisacrylamide (DHEBA), 0.25% TEMED, and 0.25% ammonium persulfate (APS). Cells were embedded by covering them with the gel mixture, allowing polymerization at room temperature for 15 minutes, followed by 1.5 hours in a humidified 37°C incubator.

The hydrogel was denatured in a solution containing 200 mM SDS, 100 mM NaCl, and 50 mM Tris (pH 6.8) for 15 minutes at room temperature, followed by incubation at 73°C for 1 hour. After denaturation, the hydrogel was transferred to a dish containing deionized water for the expansion phase. Water was replaced twice, with 1-hour intervals, and then the gel was allowed to expand overnight at room temperature. The final hydrogel diameter reached approximately 4 cm, ensuring optimal sample expansion.

The following imaging acquisitions were performed using the Dragonfly confocal imaging system. Gel samples were imaged with a 60× oil-immersion objective (1.6 NA). Multi-channel images were acquired sequentially. The resulting TIFF images were analyzed using custom-developed code, previously applied to STED imaging, to identify and quantify the distribution of PCNA and RD.

### STORM Imaging and Data Analysis

The STORM imaging and analysis were adapted from our previous work^33^. HeLa cells were seeded onto glass-bottom dishes and synchronized as previously described, followed by click chemistry for EdU with dye AF647, and immunofluorescence for BrdU with dye Cy3B.

STORM imaging was conducted using a custom-built inverted microscope (IX83, Olympus) configured for wide-field excitation. The system was equipped with a 100×, 1.49 NA oil-immersion objective lens (U APO N, Olympus). A multiband dichroic mirror (Chroma) was used to reflect the excitation lasers while allowing the transmission of emitted fluorescence. Additional emission filters (Chroma) were applied to isolate fluorescence signals from individual channels. Images were acquired on an EMCCD camera (Ixon Ultra EMCCD, Andor) with a resolution of 512*512 pixels (115 nm per pixel). A near-infrared laser was utilized to maintain precise focus during acquisition.

TetraSpeck Microspheres (0.1 μm in diameter, Thermo, T7279) were incubated with the sample at a 1:50000 ratio for 1 h, followed by three washes with PBS to remove non-adherent beads. Imaging was performed in a buffer composed of 10% (w/v) glucose, 20 mM NaCl, 100 mM Tris-HCl, 600 μg/mL glucose oxidase, 60 μg/mL catalase, and 1% (v/v) β-mercaptoethanol. Fluorophores were activated with a 405-nm laser (OBIS, Coherent) and excited using 561-nm and 647-nm lasers (MBP Communications), depending on the channel. The laser power densities at the sample plane were 1 kW/cm² for the 647 nm laser and 2 kW/cm² for the 561 nm laser.

STORM imaging data were acquired with a 30-ms exposure time per frame, generating a total of 60,000 frames for each channel (561 nm and 647 nm). After imaging, the STORM buffer was removed by washing the samples three times with PBS, and samples were stored in PBS at 4 °C shortly.

For the analysis of single-molecule localizations, the target cells and TetraSpeck Microspheres were first manually selected from the acquired images. Channel correction and drift correction were carried out using ThunderSTORM^91^ by aligning the positions of the microspheres. Following drift correction, single-molecule reconstructions and channel filtering were performed independently for the 647 nm and 561 nm channels using ThunderSTORM.

For spatial clustering, SR-Tesseler^92^ was employed for threshold-based segmentation of the processed images. A Voronoi diagram was constructed for each channel, with the 561 nm channel used to define RDs and the 647 nm channel for replication origins. Clustering was performed by applying appropriate density factors, with a density factor of 0.5 for defining RDs in the 561 nm channel, and a density factor of 5 for identifying origins in the 647 nm channel. The identified clusters were analyzed by calculating key spatial features, including the number of clusters and diameter of each cluster.

## Statistical Analysis

Statistical analyses were conducted using GraphPad Prism software (version 9). Unless otherwise specified, an unpaired two-sample parametric t-test was utilized to assess significance levels, denoted as: ****P < 0.0001, ***P < 0.0005, **P < 0.01, *P < 0.05; ns, not significant. In scatter plots, horizontal lines represent mean values, with error bars indicating ± standard deviation (s.d.). For violin or box plots, the central line represents the median, while the upper and lower bounds of the box indicate the 25th and 75th percentiles, respectively. Whiskers extend to data points within 1.5× the interquartile range of the box limits.

## Data availability

The authors declare that the data supporting the findings of this study are available within the paper and its supplementary information files. The high-throughput sequencing data have been uploaded to the GEO database with accession numbers (GSE289473, GSE289474, GSE289475). Additional analysis data and code can be obtained by contacting the corresponding author.

## Code availability

The computation details are described in the Methods section. The related codes are provided on the website.

## Acknowledgements

We thank Dr. Huiwen Hao for her valuable suggestions on imaging experiments. We are grateful to Dr. Wu Honggui from the Xiaoliang Xie Laboratory for technical advice on BrdU-seq experiments. We also acknowledge the Fenghuang Imaging Platform for providing instrument support.

## Author contributions

M.Z., Y.S., and Y.L. conceptualized the research problem. M.Z. and Y.S. designed the experiments. M.Z. completed all super-resolution imaging, PCNA single-molecule tracking imaging, parts of confocal imaging, and experiments for BrdU-seq, CUT&Tag, and RNA-seq, and led the experimental data analysis.

W.Z. analyzed the sequencing data, including BrdU-seq, Hi-C, and CUT&Tag. Z.X. contributed to STORM imaging, the construction of 231 knockout cell lines, Lamin A/C complementation experiments, and imaging experiments related to DNA damage. L.C. performed the Hi-C and chromatin loci tracking experiments. X.W. provided the original code for image analysis. Y.B. carried out the DNA combing experiments. X.G. built the STORM imaging system. T.L. provided technical guidance for STED experiments and advice on dye selection. Q.L., C.L. and D.X. gave useful advice for designing the project.

Q.P.S. and Y.S. supervised the project. M.Z., Q.P.S. and Y.S. wrote the manuscript.

